# Misfolded α-synuclein causes hyperactive respiration without functional deficit in live neuroblastoma cells

**DOI:** 10.1101/663815

**Authors:** C.L Ugalde, S.J Annesley, S Gordon, K Mroczek, M.A Perugini, V.A Lawson, P.R Fisher, D.I Finkelstein, A.F Hill

**Affiliations:** La Trobe Institute for Molecular Science, La Trobe University, Bundoora, Victoria, Australia 3082; Department of Microbiology and Immunology, University of Melbourne, Parkville, Victoria, Australia 3052; Howard Florey Institute of Neuroscience and Mental Health, Parkville, Victoria, Australia 3052; Department of Biochemistry and Molecular Biology, University of Melbourne, Parkville, Victoria, Australia 3052; Department of Physiology, Anatomy and Microbiology, La Trobe University, Bundoora, Victoria, Australia 3082; La Trobe Comprehensive Proteomics Platform, La Trobe Institute for Molecular Science, La Trobe University, Bundoora, Victoria 3086, Australia

**Keywords:** α-synuclein, synucleinopathy, Parkinson’s disease, mitochondria, cardiolipin, PMCA

## Abstract

The misfolding and aggregation of the largely disordered protein, α-synuclein, is a central pathogenic event that occurs in the synucleinopathies; a group of neurodegenerative disorders that includes Parkinson’s disease. While there is a clear link between protein misfolding and neuronal vulnerability, the precise pathogenic mechanisms employed by disease-associated α-synuclein are unresolved. Here, we studied the pathogenicity of misfolded α-synuclein produced using the Protein Misfolding Cyclic Amplification (PMCA) assay. To do this, previous published methods were adapted to allow PMCA-induced protein fibrillization to occur under non-toxic conditions. Insight into potential intracellular targets of misfolded α-synuclein was obtained using an unbiased lipid screen of 15 biologically relevant lipids that identified cardiolipin (CA) as a potential binding partner for PMCA-generated misfolded α-synuclein. To investigate if such an interaction can impact the properties of α-synuclein misfolding, protein fibrillization was carried out in the presence of the lipid. We show CA both accelerates the rate of α-synuclein fibrillization and produces species that harbour enhanced resistance to proteolysis. Because CA is virtually exclusively expressed in the inner mitochondrial membrane, we then assessed the ability of these misfolded species to alter mitochondrial respiration in live non-transgenic SH-SY5Y neuroblastoma cells. Extensive analysis revealed misfolded α-synuclein causes hyperactive mitochondrial respiration without causing any functional deficit. These data give strong support for the mitochondrion as a target for misfolded α-synuclein and reveals persistent, hyperactive respiration as a potential up-stream pathogenic event associated with the synucleinopathies.

**Summary statement:** Misfolded α-synuclein that was produced using the Protein Misfolding Cyclic Amplification (PMCA) assay was found to associate with cardiolipin and cause hyperactive respiration in neuronal cells.

## Introduction

Parkinson’s disease (PD) is a slowly progressing movement disorder clinically defined by the appearance of bradykinesia, rigidity and/or tremor. Pathological features of PD are the appearance of Lewy Bodies, which are intracellular protein deposits predominately composed of misfolded α-synuclein and neuronal loss. PD is one of a group of neurodegenerative conditions called synucleinopathies that have in common the aggregation of misfolded α-synuclein. The profiles of neuronal vulnerability and protein deposition within neurons and glial cells vary amongst the conditions (Galvin et al., 2001; Surmeier et al., 2017) and PD is associated with the progressive loss of dopaminergic neurons in the substania nigra pars compacta.

Largely an unstructured protein that is localized to the synapse under normal physiological conditions, the misfolding of α-synuclein is strongly linked to the development of disease. This is evidenced by the finding that various point mutations in the gene encoding α-synuclein, *SNCA* (such as A53T), causes early onset familial disease (Appel-Cresswell et al., 2013; Kruger et al., 1998; Lesage et al., 2013; Polymeropoulos et al., 1997; Proukakis et al., 2013; Zarranz et al., 2004) as do gene duplications (Chartier-Harlin et al., 2004) and triplications (Singleton et al., 2003). Likewise, there is also now a large body of data showing the ability of α-synuclein to adopt pathogenic properties upon misfolding (reviewed in: (Ugalde et al., 2016)).

Despite the numerous data describing the pathogenicity of misfolded α-synuclein, the precise molecular mechanisms α-synuclein employs that contribute to the pathogenesis of the synucleinopathies is currently unclear. A consideration to studying these disorders is the complex nature of α-synuclein misfolding, which as an amyloid forming protein, can adopt numerous misfolded conformations. Precursors to the mature fibrils abundant in LBs, small soluble oligomers and protofibrils also present in the brain in PD (Garcia-Esparcia et al., 2015; Sharon et al., 2003; Tofaris et al., 2003) and critically, oligomers are considered to be the most toxic conformation (reviewed in (Ugalde et al., 2016)). The process of α-synuclein fibril-formation sees these highly dynamic species exist as part of a heterogeneous mix of various sized conformations that maintain a state of equilibrium and through interspecies interactions, can both expand and contract to higher or lower ordered structures (Cremades et al., 2012; Roberts and Brown, 2015).

Many studies examining the properties of pathogenic α-synuclein induce misfolding in bacterially-expressed recombinant protein to produce a homogenous population of a defined misfolded structure. While these common techniques of fibrillization are useful to attribute pathogenic properties to a specific conformation, they are not reflective of the numerous conformations of pathogenic α-synuclein that exist *in vivo*. A novel fibrillization technique that may alleviate these issues is Protein Misfolding Cyclic Amplification (PMCA). PMCA was originally developed to study the generation of prions (Saborio et al., 2001), which are proteinaceous and transmissible neurodegenerative agents primarily composed of PrP^Sc^ that is a misfolded form of the normal protein, PrP^C^. Recently, the PMCA has been shown capable of producing misfolded forms of α-synuclein that appear to encompass various conformations, including oligomers (Herva et al., 2014). Current studies on the pathogenicity of α-synuclein species formed via PMCA are limited to low concentrations of dilute protein due to a high concentration of detergent in the conversion buffer. As such, we sought to study the properties of PMCA-generated α-synuclein by adapting published methods to produce misfolded species under non-toxic conditions and extensively characterizing formed species. Next, to gain insight into the pathogenic properties of misfolded α-synuclein, an unbiased lipid screen of 15 biologically relevant lipids was performed. We show that PMCA-generated α-synuclein associates with cardiolipin (CA), and further assessment reveals this lipid can both accelerate fibrillization and modify the biophysical and biochemical properties of α-synuclein species formed by enhancing their resistance to proteolysis and increasing insoluble load. Given that CA is a lipid almost exclusively expressed in the inner mitochondrial membrane (IMM), we then sought to determine if this interaction has relevance to cellular functioning by assessing the ability of PMCA-generated α-synuclein to modulate mitochondrial respiration in live non-transgenic neuroblastoma cells. Comprehensive analysis using the Seahorse XF Analyzer showed misfolded α-synuclein causes hyperactive mitochondrial respiration without causing dysfunctional activity at any complex. This also occurred independently of upstream metabolic changes in glycolysis. This rigorous data in cells with normal endogenous protein suggests the mitochondrion is a target for misfolded α-synuclein and that hyperactive respiration may be an early, up-stream pathogenic process neuronal cells experience in the synucleinopathy disorders.

## Results

### PMCA can produce misfolded α-synuclein species in prion conversion buffer (PCB)

Generating misfolded α-synuclein species using PMCA was first developed following the protocol outlined by Herva et al. (Herva et al., 2014) (Figure 1A). Because the buffer used by Herva et al. to dissolve α-synuclein is routinely used as the conversion buffer for prion PMCA reactions, it is hereafter referred to as the prion conversion buffer (PCB). The extent of fibrillization of α-synuclein in PCB following exposure to PMCA for 24 h was assessed using western immunoblot, transmission electron microscopy (TEM) and thioflavin T (ThT) fluorescence. Here, western immunoblot was employed to identify small conformations of misfolded α-synuclein that may be resolved by SDS-PAGE and showed samples subjected to PMCA contained a range of various sized species, with monomeric protein observed at 14 kDa and dimer, trimer and tetramer oligomers identified by their increase in molecular weight by intervals of the weight of the monomer (Figure 1B). Species greater than a tetramer, separated by the size of one monomeric unit were identified all the way up to the high molecular weight range of the gel. In comparison, a non-PMCA sample stored at −80°C during the PMCA process was almost entirely composed of monomeric protein (Figure 1B). TEM was next used to confirm the presence of large mature α-synuclein fibrils. Consistent with the TEM images reported by Herva et al. (Herva et al., 2014), PMCA generated α-synuclein fibrils were found to be rod-like in structure. No aggregates were observed in the −80°C sample (Figure 1C). The compound ThT exhibits fluorescence upon binding to β-sheet structures (Biancalana and Koide, 2010; Vassar and Culling, 1959), and hence is universally used to detect amyloid. Upon exposure to 24 h PMCA, α-synuclein adopted strong ThT reactivity, while the monomeric −80°C sample displayed no fluorescent signal (Figure 1D). Collectively these data confirm PMCA is a useful tool to produce misfolded α-synuclein.

**Figure 1:**
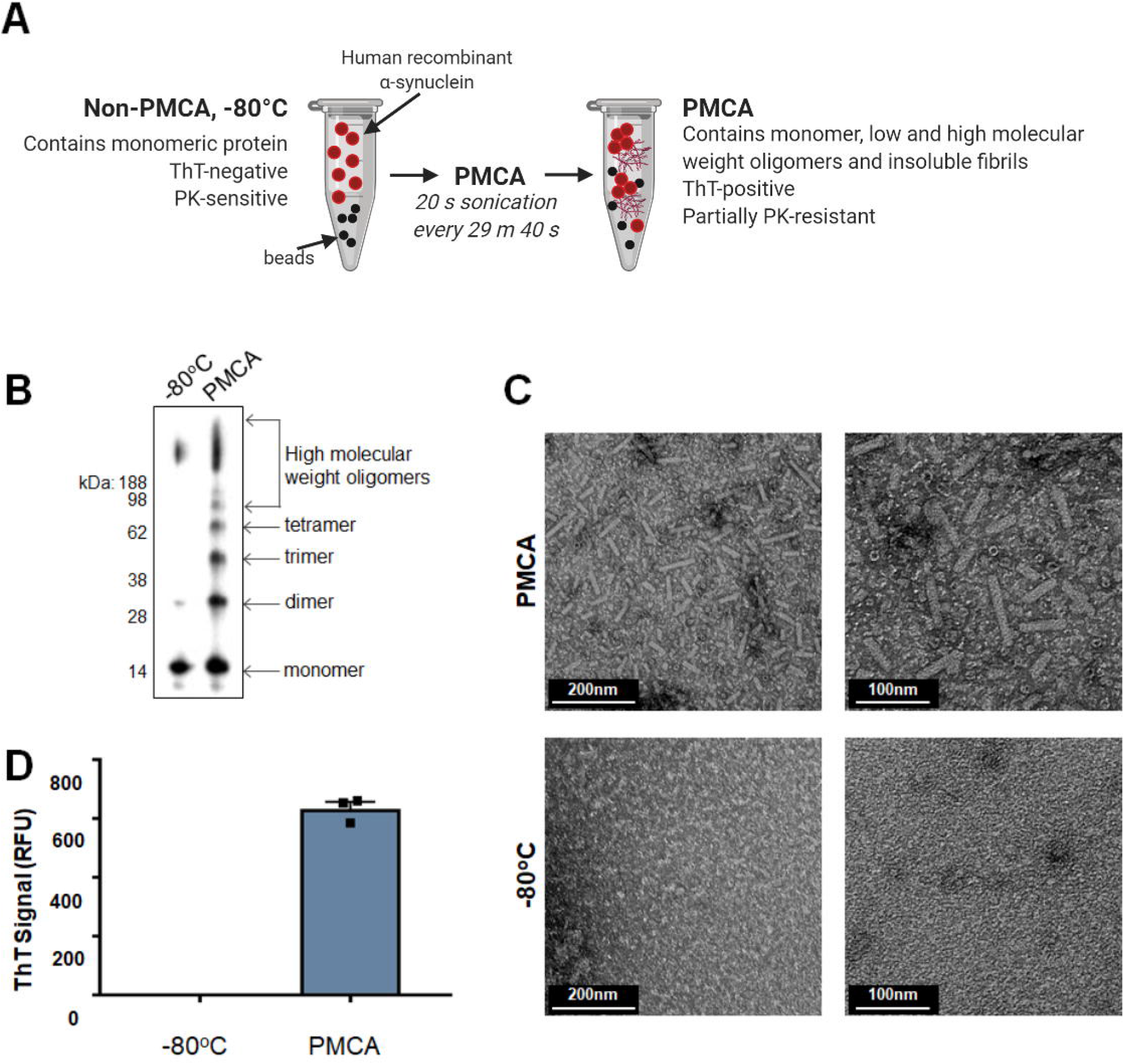
α-synuclein PMCA. (A) Workflow of αsyn PMCA: Lyophilized recombinant wild-type protein was reconstituted in buffer to 90 μM and 60 μL aliquoted into PCR tubes. Samples were exposed to PMCA in the presence of beads. Sonications were set to occur for 20 sec every 29 min 40 sec for a total process time of 72 h. Extent of fibrillization in protein exposed to PMCA and non-PMCA controls (−80°C) was assessed by (B) western immunoblot analysis using α-synuclein-specific monoclonal antibody MJFR1 (amino acid specificity: 118-123) (C) transmission electron microscopy (representative images) and (D) ThT fluorescence, where the values obtained for each experimental replicate represented the average fluorescence of triplicate wells after subtraction of a blank well to account for background fluorescence (RFU= relative fluorescent units). Data presented as mean±SEM (n=3) and represent every experimental replicate performed.

### PMCA-induced misfolding of α-synuclein can occur under non-toxic conditions

Because the PCB used to reconstitute protein in Figure 1 contains Triton X-100, a limitation to studying the pathogenicity of PMCA-generated α-synuclein is the inherent toxicity of the buffer. Accordingly, in order to progress to studies on the pathogenicity of PMCA-generated α-synuclein, it was necessary to isolate these species in a buffer that did not contain detergent. Given the small size and highly dynamic nature of oligomers, buffer exchange or centrifugation were not considered viable options to collect or maintain the integrity of small, oligomeric species that may be produced in this system. Instead, the replacement of the conversion buffer, PCB, with a non-toxic analogue was pursued. In order to address this, the extent of PMCA-induced fibrillization of α-synuclein was assessed following its reconstitution in several buffers assumed to be non-toxic. The following buffers were tested: PBS, PBS+150 mM NaCl (PBSN), TBS and TBS+150 mM NaCl (TBSN). These samples were exposed to 24 h PMCA as per the same conditions detailed in Figure 1 and were compared to an equivalent sample of α-synuclein dissolved in PCB buffer (α-synuclein:PCB). Western immunoblot and ThT were used as readouts to compare the extent of α-synuclein fibrillization in the various buffers (Supplementary Figure 1). The ThT data showed that following PMCA, all buffers were able to produce species that contained ThT-reactive conformations (Supplementary Figure 1A). Resolving samples on SDS-PAGE and immunodetection of α-synuclein gave further information on the α-synuclein species produced by the buffers (Supplementary Figure 1B). This assay revealed differences in the efficiency of the buffers to induce fibrillization, with PCB clearly being the most proficient buffer for producing oligomeric species. The degree of fibrillization was variable amongst the non-toxic buffers with PBSN being the most effective non-toxic buffer for α-synuclein misfolding. This observation is consistent with studies showing salt concentrations are positively associated with α-synuclein fibrillization (Hoyer et al., 2002).

### Characterisation of PMCA-generated α-synuclein misfolding in PBSN

Given that none of the non-toxic buffers were as effective as PCB in facilitating misfolding of α-synuclein based on western immunoblot, the protocol was further adapted with the most efficient nontoxic buffer, PBSN (Figure 2). To assess this, the total process time of 24 h was extended to determine if PBSN may produce comparable misfolded α-synuclein content to PCB if exposed to PMCA for a longer period. To this end, an incremental time-course experiment was performed where α-synuclein:PBSN was subjected to PMCA for a total process time of: 0, 24, 48 and 72 h. These samples were compared to α-synuclein:PCB exposed to 24 h PMCA. Western immunoblot analysis revealed the abundance of misfolded species produced in PBSN continues to amplify beyond a 24 h PMCA process time, with time-dependent elevations in misfolded content observed in samples exposed to PMCA for 48 h and 72 h. Following exposure to PMCA for 72 h, the presence of oligomeric species in α-synuclein:PBSN was comparable to 24 h α-synuclein:PCB (Figure 2A). Because the use of western immunoblot to detect misfolded α-synuclein may be compromised by artefacts associated with antibody epitope recognition, silver stain was also employed to detect total protein. Using this method confirmed the PMCA induces the generation of α-synuclein oligomers that are time dependent (Figure 2B). However, unlike the western immunoblot data (Figure 1A), saturation was observed in the monomeric band (Figure 1B; indicated by an asterisk). Collectively, these data show that while PMCA is capable of producing α-synuclein oligomers, the unsaturated signals from the western immunoblot may not represent the exact ratios of species that resolve through SDS-PAGE, likely related to antibody epitope specificity.

**Figure 2:**
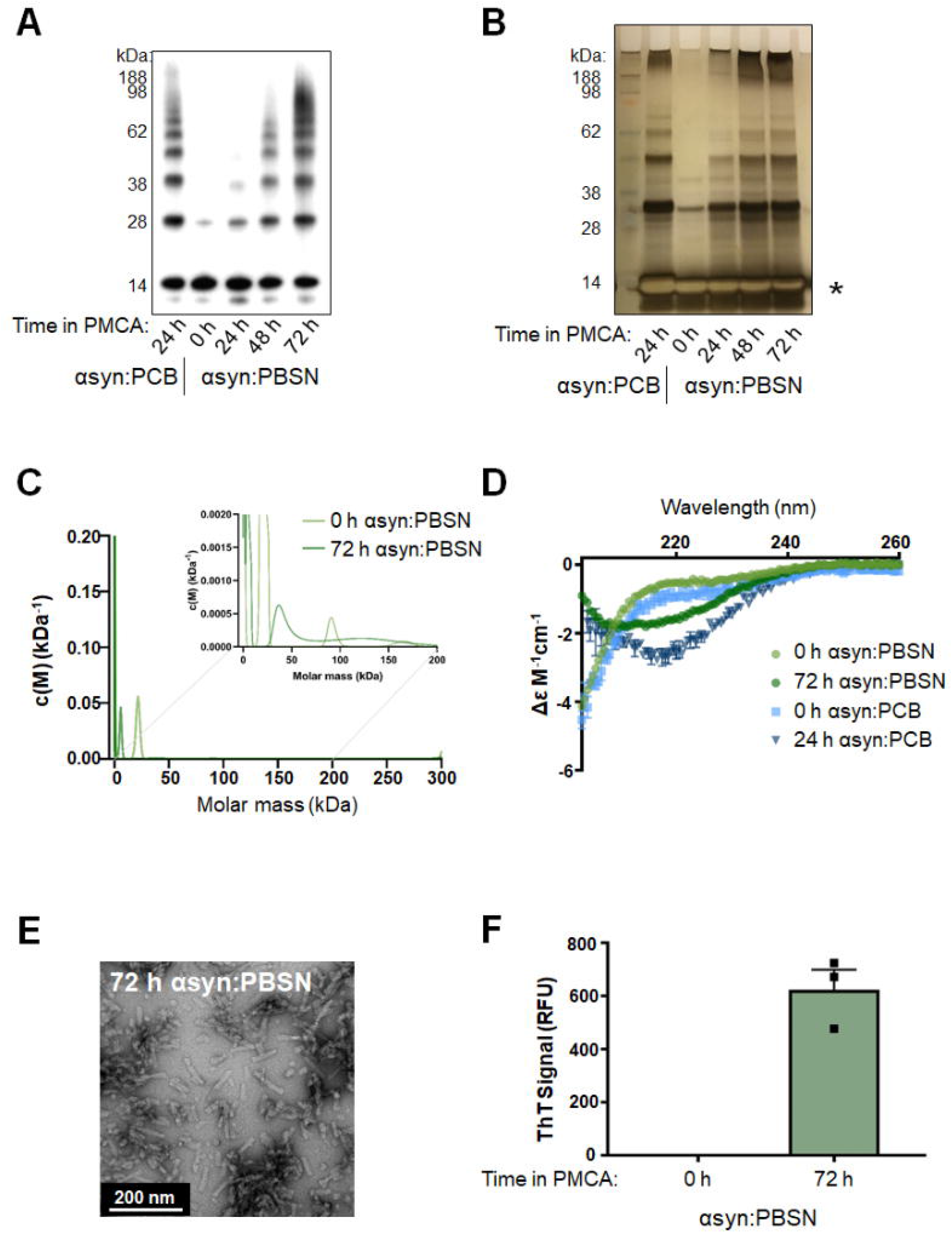
Production of misfolded α-synuclein species in PBSN buffer produces comparable misfolded content to α-synuclein species generated in prion conversion buffer (PCB) Lyophilized recombinant wild-type protein was reconstituted in PBSN (α-synuclein:PBSN) or PCB (α-synuclein:PCB) and subjected to PMCA for a total process time of 0, 24, 48 or 72 h. (A) Extent of fibrillization in samples was assessed using western immunoblot analysis using α-synuclein-specific monoclonal antibody MJFR1 (amino acid specificity: 118-123) and (B) silver stain. (C) Analytical ultracentrifugation sedimentation velocity analyses of non-PMCA and PMCA-generated α-synuclein. Continuous mass distribution (c(M)) distributions are shown as a function of molar mass. Insert shows zoomed in image of values between 0 – 200 kDa. Best fits to experimental data yielded an RMSD of 0.005 461 and 0.004 502, ff0 of 2.2 and 1.1, and Runs Test Z of 12.23 and 0.59 for non-PMCA and PMCA-generated α-synuclein, respectively. (D) Secondary structure of non-PMCA (0 h) or PMCA-generated (24 or 72 h) α-synuclein:PBSN and α-synuclein:PCB was determined using circular dichroism (CD) spectroscopy (n=1, representative spectra). (E) Transmission electron microscopy of fibrils produced in PBSN after 72 h PMCA (representative image). (F) α-synuclein:PBSN exposed to PMCA for 72 h or non-PMCA (0 h) monomeric controls were further characterized by ThT fluorescence, where the values obtained for each experimental replicate represented the average fluorescence of triplicate wells after subtraction of a blank well to account for background fluorescence (RFU= relative fluorescent units). Data presented as mean±SEM (n=3) and represent every experimental replicate performed.

Following on from these findings, sedimentation velocity experiments were next conducted in the analytical ultracentrifuge (AUC) to provide an in-solution assessment of the presence of oligomeric species in PMCA-generated α-synuclein. In order to align with the experimental procedure of other assays, these experiments were conducted at 25°C and 37°C for non-PMCA (0 h α-synuclein:PBSN) and PMCA-generated α-synuclein (72 h α-synuclein:PBSN), respectively. Continuous mass distribution profiles (c(M)) were given for both proteins by fitting sedimentation profiles to the c(M) distribution model (Schuck, 2000; Schuck et al., 2002). As shown in Figure 2C, both distributions show a single predominant peak of mass 6–20 kDa that is a reasonable approximation of the α-synuclein monomer (≈14 kDa). A best-fit, weight-average f/f_0_ value of 2.2 was found for the non-PMCA α-synuclein sample, indicating a very prolate solution conformation, while contrastingly the f/f_0_ for PMCA-treated α-synuclein was 1.1, consistent with a more globular shape. Closer inspection of the region spanning 0 kDa to 200 kDa (insert, Figure 2C) reveals the presence of several broad peaks between 20 – 60 and 100 – 150. Compellingly, the molecular assignments for these peaks are in good general agreement with the western immunoblot and silver stain data for the equivalent samples. Residuals for fits for AUC data are provided in Supplementary Figure 2. Thus, these data support the presence of low molecular weight oligomers in solution in PMCA-generated α-synuclein:PBSN.

For the AUC data acquisition, a higher rotor speed of 42,000 was selected to maximise resolution of low MW components. To assess the extent to which depletion of high sedimentation coefficient solutes (e.g. fibrils) occurs, we compared absorbance-detected sedimentation profiles used for c(M) distribution models (42,000 rpm) with those collected at a lower rotor speed (3000 rpm), where sedimentation of these larger components will be minimal. An approximate sixfold loss of absorbance intensity is apparent after acceleration to the higher rotor speed of 42 000 rpm, indicating rapid sedimentation of solutes with high sedimentation coefficients. This is further support of an expansion in fibrillar content with PMCA treatment. Contrastingly, under these conditions we find negligible loss of absorbance signal for non-PMCA α-synuclein.

Having characterised misfolded protein using AUC, further assessment was conducted on fibrillar species formed in α-synuclein:PBSN. Circular dichroism (CD) spectra was obtained for non-PMCA (0 h) and PMCA-generated α-synuclein in both PCB and PBSN. Consistent with the organized rearrangement of α-synuclein upon misfolding, non-PMCA samples of both buffers showed largely random coil present that shifted to become rich in β-sheet following PMCA (Figure 2D). Likewise, using TEM, the morphology of fibrillar species produced in PBSN after PMCA for 72 h (Figure 2E), was found to be consistent with those produced in PCB (Figure 1C and Herva et al. (Herva et al., 2014)). Finally, similar to 24 h α-synuclein:PCB shown in Figure 1D, 72 h α-synuclein:PBSN was ThT-reactive (Figure 2F). In conclusion, while it cannot be presumed PBSN precisely mirrors the mechanism of misfolding that occurs using PCB (for example, due to the effect of triton-X 100 that is present in PCB), collectively these data show 72 h is an appropriate PMCA process time to produce misfolded species in PBSN that share many similar amyloidogenic and oligomeric properties as protein produced in PCB.

A necessary component to confirming that PBSN is a suitable buffer to produce misfolded α-synuclein that can facilitate studies on its pathogenicity was to confirm PBSN is non-toxic when applied to cultured cells. To address this, the ability of the buffer alone (void of α-synuclein protein) to cause cell death in neuroblastoma SH-SY5Y cells was measured via the release of lactate dehydrogenase (LDH) into conditioned media. In order to test the usefulness of PBSN-based buffers in PD-model systems, LDH assays were also performed using the A4 cell line that are SH-SY5Y cells stably transfected to overexpress wild-type α-synuclein. Prior to LDH measurements, confirmation of elevated α-synuclein expression in A4 cells compared to untransfected SH-SY5Y counterparts came from western immunoblot and immunofluorescence (Supplementary Figure 3A-B). Both cell lines were treated with PCB or PBSN such that the total volume of culture media was composed of 0.5, 1, 5 or 10% v/v buffer and LDH measured 24 h later (Supplementary Figure 3C,E). Results showed a clear dose-dependent increase in toxicity of PCB (buffer only), with 10% PCB eliciting a near-100% cell death response for SH-SY5Y and A4 cells. In contrast, PBSN (buffer only) showed negligible toxicity at all concentrations tested in both cell lines. The cytotoxicity of the buffers measured by LDH was confirmed by differential interference contrast (DIC) images of cells exposed to PCB or PBSN (Supplementary Figure 3D,F).

Collectively, these experiments confirm PBSN is a suitable buffer to produce α-synuclein species under non-toxic conditions, which greatly improves the capacity of the PMCA to be used as a tool to study pathogenic properties of misfolded α-synuclein species. As such, all subsequent experiments using PMCA-generated misfolded α-synuclein were produced using this optimized method.

### PMCA-generated misfolded α-synuclein selectively associates with cardiolipin

The development of a method to produce misfolded α-synuclein under non-toxic conditions using PMCA allowed the commencement of the next aim, which was to identify potential pathogenic mechanisms of misfolded α-synuclein. While numerous mechanisms of how α-synuclein causes cellular damage have been reported, a common theme unifying several of these mechanisms appears to involve the interaction of misfolded α-synuclein with lipid species (reviewed in: (Ugalde et al., 2019)). Therefore, to gain insight on potential mechanisms to pursue, the ability of misfolded α-synuclein to interact with various lipids was assessed using a hydrophobic membrane that had been dotted with 15 long chain (>diC16) highly pure synthetic lipid analogues and a blank control. This membrane was incubated with misfolded (PMCA) or monomeric (−80°C) α-synuclein and the degree of protein binding to the lipids assessed via immunoblot using the α-synuclein-specific antibody MJFR1. These qualitative results revealed that, compared to the −80°C sample, PMCA α-synuclein exhibited a general elevation in both lipid-specific and non-specific binding to the membrane (Figure 3A-B). By far, α-synuclein exhibited the strongest affinity to cardiolipin (CA). Interestingly, the interaction of α-synuclein with these lipid species was conformation specific, with no reactivity with any lipid observed after incubation with the −80°C sample (Figure 3B). Given that this sample contains monomeric protein and no ordered aggregates, it can be inferred that the protein interacting with lipids in the PMCA sample are ordered misfolded structures.

**Figure 3:**
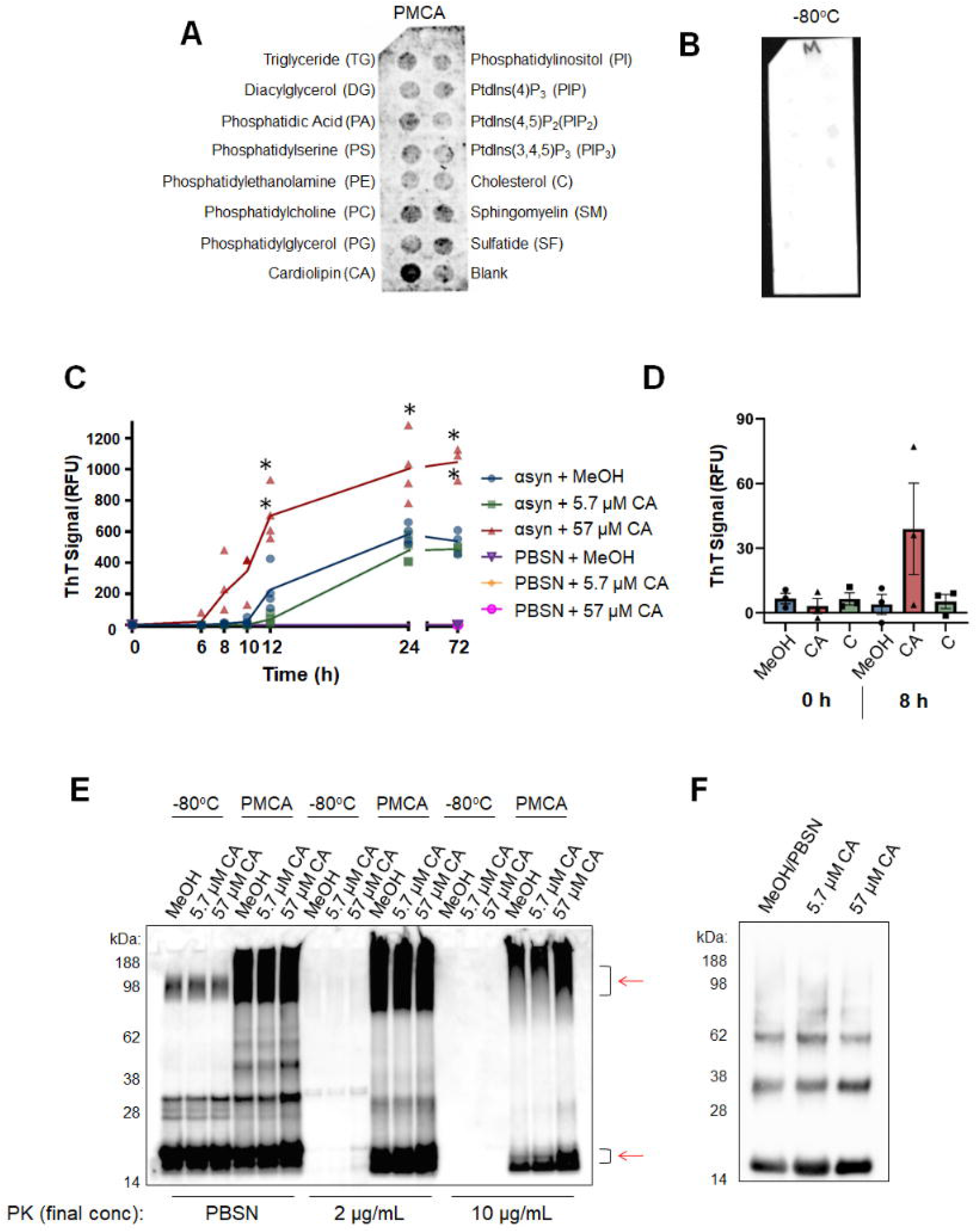
PMCA-generated α-synuclein selectively associates with cardiolipin (CA). A hydrophobic membrane strip dotted with 15 different lipids (long-chain >diC16 synthetic analogues) was incubated with PMCA-generated α-synuclein (A) Binding of α-synuclein to lipids was assessed via western immunoblot using α-synuclein-specific monoclonal antibody MJFR1 (amino acid specificity: 118-123). (B) Unlike PMCA-generated misfolded α-synuclein, monomeric α-synuclein (−80°C) did not show affinity to any lipid class, n=1. To determine if cardiolipin (CA) modulates α-synuclein fibrillization, CA was dissolved in MeOH/PBSN (MeOH) and added to wild-type α-synuclein for a final concentration of 5.7 and 57 μM CA. MeOH/PBSN alone added to α-synuclein: PBSN served to control for the effect of the buffer. (C) Samples were subjected to PMCA for a total process time of 0, 6, 8, 10, 12, 24 and 72 h. Tubes containing PBSN only (no α-synuclein) with buffer or CA exposed to PMCA for 0 and 72 h served to detect any inherent fluorescence caused by CA or the buffer. Extent of fibrillization was assessed by measuring ThT fluorescence, where the values obtained for each experimental replicate represented the average fluorescence of triplicate wells after subtraction of a blank well to account for background fluorescence (RFU= relative fluorescent units). Data presented as mean±SEM (n=4 for all groups except for 0 h and 72 h which were n=3) and represent every experimental replicate performed. To determine the effect of CA on α-synuclein fibrillization compared to control samples, a mixed-effects ANOVA with Dunnett’s multiple comparisons test was employed *p<0.05, **p<0.01. (D) ThT was repeated in the presence of CA or cholesterol (C) which showed no binding affinity to α-synuclein in panel A. ThT was measured after 8 h which represented the earliest detectable rise in ThT by CA observed in panel C. The values obtained for each experimental replicate represented the average fluorescence of triplicate wells after subtraction of a blank well to account for background fluorescence. Data presented as mean±SEM (n=3) and represent every experimental replicate performed. (E) The biochemical properties of α-synuclein species formed after 72 h PMCA (PMCA) or not (−80°C) was assessed by its resistance to proteolysis following digestion in proteinase K (PK) at final concentration of 0, 2 or 10 μg/mL. PK-resistant species were detected by immunoblot using MJFR1. Areas of differential PK resistance between PMCA-generated species are identified by red arrows. n=1. (F) Ultracentrifugation was performed on α-synuclein in the presence of 5.7 and 57 μM CA or buffer control to determine the insoluble load. n=1

The identification of CA as a lipid class with high binding affinity to misfolded α-synuclein using an unbiased screen merited further analysis into whether this lipid can modulate α-synuclein fibrillization. To do this, ThT was employed to study the kinetics of α-synuclein fibrillization in the presence of the lipid. After reconstitution in methanol (MeOH) and PBSN, CA was added to α-synuclein:PBSN for a final concentration of 57 μM and 5.7 μM. MeOH alone was added to tubes with equivalent protein to serve as a control for the effect of the buffer. The ability of CA to modulate α-synuclein fibrillization was determined by measuring ThT fluoresence in individual tubes containing CA exposed to PMCA for a total process time of: 0, 6, 8, 10, 12, 24 or 72 h and analysed compared to equivalent CA-free control samples. Here, compared to tubes containing α-synuclein with MeOH (buffer only), the addition of 57 μM CA accelerated α-synuclein fibrillization which was evidenced by a statistically significant elevation in ThT first identified after 10 h (p=0.031) (Figure 3C). This elevation was sustained in all subsequent time-points (12, 24 and 72 h; p=0.009, 0.041 and 0.006, respectively). No significant differences were identified between the control group and the lower concentration of CA (5.7 μM CA). Importantly, neither the lipid nor MeOH exhibited any florescence signal in the absence of α-synuclein (Figure 3C). In order to determine whether the ability of CA to accelerate α-synuclein fibrillization was confounded by molecular crowding of the lipid, ThT was performed again using CA alongside a lipid found to not associate with the protein as determined by the lipid strip (Figure 3A). Here, CA or the equivalent molar concentration (57 μM) of cholesterol (C) was added to α-synuclein and ThT measured after 8 h exposure to PMCA. This time-point was chosen to represent the earliest CA-induced elevation in ThT (Figure 3C) and confirmed that, at this timepoint, only CA (and not C or buffer controls) exhibited PMCA-induced elevations in ThT (Figure 3D). No ThT reactivity was observed in any sample at 0 h.

Having observed the ability of CA to modulate α-synuclein fibrillization detected using a fluorescent technique, we sought to complement these observations by examining what effect this has on the misfolded protein produced. It has been shown that species formed by PMCA are more resistant to proteolysis (Herva et al., 2014) which may be a useful assay to detect changes in the population of species formed. Accordingly, the effect of CA on the misfolding of α-synuclein was next assessed by exposing PMCA products to proteinase K (PK) digestion and performing western immunoblot. PMCA of α-synuclein was performed in reactions containing CA (as per the conditions for Figure 3CD) or buffer for 72 h prior to being exposed to 0, 2 or 10 μg/mL PK. These samples were compared to corresponding tubes that did not undergo PMCA (−80°C). Results showed in the absence of proteolysis or low concentrations of PK (2 μg/mL), there appeared to be more oligomeric content formed when α-synuclein was incubated with 57 μM CA. Upon treatment with 10 μg/mL PK, an increase in both high and low molecular weight protein was observed in the PMCA sample containing 57 μM CA, compared to all other groups (Figure 3E). Densitometric analysis revealed, in the PMCA samples digested with 10 μg/mL PK, the total protein abundance in the sample containing 57 μM CA was 1.38 fold higher than its buffer control (MeOH/PBSN) equivalent. This altered expression appeared to be due to changes in PK resistant material in the molecular weight range of 90 – 100 kDa and 15 – 20 kDa. On the western immunoblot in Figure 3E, these regions are identified by red arrows. Given that elevated reactivity was observed in both the high and low molecular weight region, these two populations of PK resistant protein likely reflect the large species that were able to resolve further through the gel following PK digestion and their concomitant cleavage products. Both 2 and 10 μg/mL PK digested all protein in all −80°C groups indicating neither the CA nor buffer interfered with the activity of PK in this experiment. This assay compliments the findings of Figure 3C and indicates that the acceleration of fibrillization seen is associated with changes to the biochemical properties of formed species in this system. Given that the enhanced proteolysis resistance is potentially related to an increase in fibrillar content, PMCA-generated protein formed in the presence of MeOH (buffer control), 5.7 μM or 57 μM CA was exposed to ultracentrifugation to isolate insoluble protein load. These results showed that CA had a mild influence on increasing insoluble content whereby, compared to buffer control, protein produced in the presence of 5.7 μM and 57 μM CA had an increase in insoluble content of 11 and 13%, respectively. Collectively these studies show PMCA-generated α-synuclein associates with CA which can influence both the rate of fibrillization and the biochemical and biophysical properties of the species formed.

### PMCA-generated misfolded α-synuclein causes mitochondrial hyperactivity without any functional deficit in naïve SH-SY5Y cells

Within a cell, the expression of CA is almost exclusively localized to the inner mitochondrial membrane (IMM). Hence, the finding that misfolded α-synuclein binds selectively to CA suggests that the mitochondrion is a target for misfolded α-synuclein. A major biochemical process driven by mitochondria is oxidative phosphorylation that produces the majority of the cell’s energy in the form of adenosine triphosphate (ATP). This respiration occurs via the electron transport chain (ETC), which is a series of reactions that take place within the IMM and involves four complexes (I-IV) that collectively act to shuttle electrons between them. This process releases protons into the intermembrane space (IMS); creating a proton gradient between the IMS and the mitochondrial matrix (MM) that in turn drives the generation of ATP through the large multi-subunit enzyme complex, ATP synthase.

We sought to assess the effect misfolded α-synuclein has on mitochondrial respiration by measuring respiratory flux rates, which are the rates of electron transfer to molecular oxygen via the electron transport chain. This is performed in live cells following the incremental addition of compounds that selectively inhibit components of oxidative phosphorylation. In this regard, information of the functioning of each of the components of oxidative phosphorylation may be obtained. Unlike standard techniques that quantify products of respiration at a single time-point, the ability to measure flux changes in mitochondria is a powerful way to identify both overt and subtle alterations to respiration.

The ability of misfolded α-synuclein to alter mitochondrial respiration was measured in naïve SH-SY5Y neuroblastoma cells using the Seahorse XFe24 Analyzer. In this system, respiration is measured via changes in oxygen (referred to as oxygen consumption rate; OCR) in the media in wells containing adhered cells. This is performed using a solid-state sensor probe that detects changes in dissolved oxygen in a portion of the media over time, with multiple readings taken to increase the sensitivity of the assay. Baseline oxygen consumption rate (OCR) readings were taken on adhered cells, prior to the sequential addition of the pharmacological chemicals oligomycin, carbonyl cyanide m-chlorophenyl hydrazone (CCCP), rotenone and antimycin A. Respectively, these compounds inhibit ATP-synthase, permeabilize mitochondrial membranes to protons, inhibit complex I and complex III (in the presence of rotenone this inhibits the flow of electrons from Complex II to oxygen). The data obtained from such measurements allows the following parameters to be measured in a single experiment: basal respiration, ATP synthesis, maximum (max) respiration, Complex I activity, Complex II activity, non-respiratory, proton leak, and spare capacity. A schematic showing a typical respirometry experiment is shown in Supplementary Figure 4A.

In addition to cells exposed to misfolded α-synuclein, other treatment groups included cells incubated with monomeric α-synuclein (−80°C), buffer alone (PBSN) and cells left untreated (Untreat). The final concentration used for all treatment groups was 5% v/v, which in order to aid the delivery of protein into the cell, were co-treated with lipofectamine. Supplementary Figure 4B is an example experiment performed in the current study. In these experiments, two treatment groups (PBSN-treated and untreated) were used as controls and hence we first examined whether PBSN alone was causing any inherent changes to mitochondrial respiration compared to untreated cells. Shown in Supplementary Figure 4C-H, assessment of mitochondrial readouts found no difference between the two controls groups. Because PBSN alone did not contribute to any mitochondrial respiration parameter, these experimental groups were pooled into one group named Control for subsequent assessment compared to monomeric (−80°C) or misfolded (PMCA) protein.

Statistical analysis on independent experimental replicates showed cells exposed to misfolded α-synuclein had an elevated basal OCR compared to both monomeric and control-treated cells (Figure 4A-F). This significant elevation of OCR in the presence of misfolded α-synuclein was maintained in individual components of mitochondrial respiration; ATP synthesis, max OCR, Complex I activity and non-respiratory (Figure 4B-D, F). Importantly, no significant difference was found between −80°C, and control groups in any readout. While Complex II activity was also elevated upon exposure to misfolded α-synuclein compared to the two other groups, this difference did not reach statistical significance in comparisons between any groups (Figure 4E). However, this observation is not unexpected given the low overall contribution of Complex II to the OCR in this cell line, relative to the accuracy and reproducibility of the assay. To summarize these data, misfolded α-synuclein causes broad-spectrum elevations to all components of respiration in mitochondria of SH-SY5Y cells. This hyperactivity was specific to misfolded α-synuclein and was not observed when cells were treated with monomeric protein or control-treated cells. Supplementary Table 1 detail all significant values from statistical analysis of cellular respiration data shown in Figure 4.

**Figure 4:**
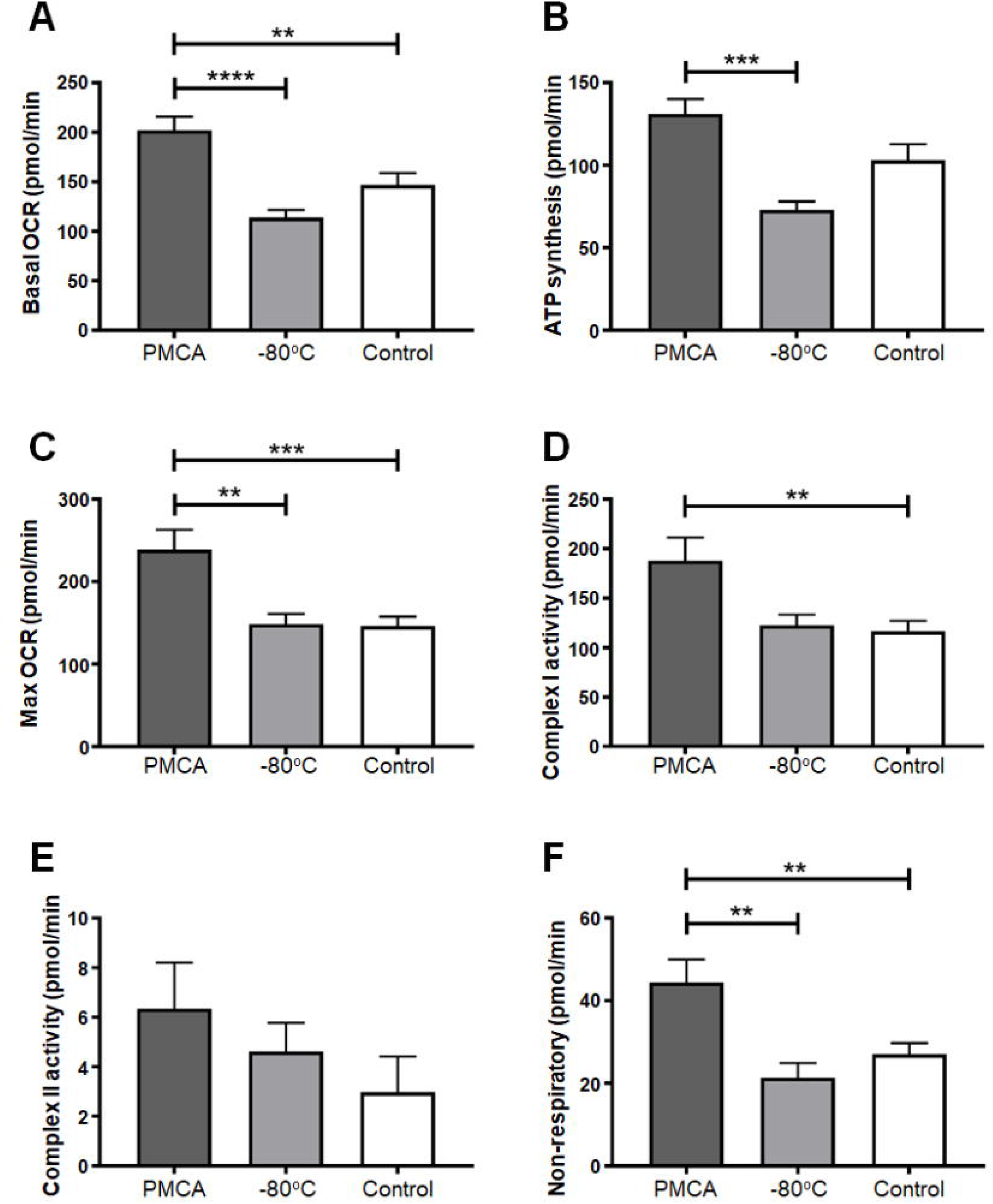
PMCA-generated misfolded α-synuclein causes mitochondrial respiration to become hyperactive in SH-SY5Y cells. SH-SY5Y cells were incubated with PMCA-generated α-synuclein, monomeric α-synuclein (−80°C) or either buffer alone or left untreated which collectively represented the Control group. Media-containing cells were plated into a well of a Seahorse XFe24 plate and mitochondria respiration measured in adhered cells using the Seahorse XF Analyzer. This was performed by detecting changes in oxygen (referred to as the oxygen consumption rate; OCR) following the addition of pharmacological agents: oligomycin, carbonyl cyanide m-chlorophenyl hydrazone (CCCP), rotenone and antimycin A. In doing so, the following parameters were measured: (C) basal OCR, (D) ATP synthesis, (E) max OCR, (F) complex I activity, (G) complex II activity and (H) non-mitochondrial respiration. The average of five wells was taken for each sample per experimental replicate. Data presented as mean±SEM (n=16, 13, 18 for groups PMCA, −80°C and Control respectively) and represent every experimental replicate performed. Statistical significance was examined by ANOVA and Tukey’s multiple comparisons test with a statistical criterion of 0.01. **p<0.01, ***p<0.001, ****p<0.0001

Further analysis of the OCR values obtained in Figure 4 was performed to determine whether the alterations in respiration identified were associated with concomitant functional deficits. The following functional readouts were calculated: basal (% max OCR), ATP (% basal), Complex I (% max OCR), non-respiratory (% basal OCR), spare capacity and proton leak. No difference was seen in any experimental group (Figure 5), indicating that misfolded α-synuclein does not affect the function and thus fractional contribution to respiration by any of the individual complexes. Therefore, while misfolded α-synuclein causes mitochondrial hyperactivity, it does not translate to any kind of observable dysfunction in the mitochondria of neuroblastoma SH-SY5Y cells.

**Figure 5:**
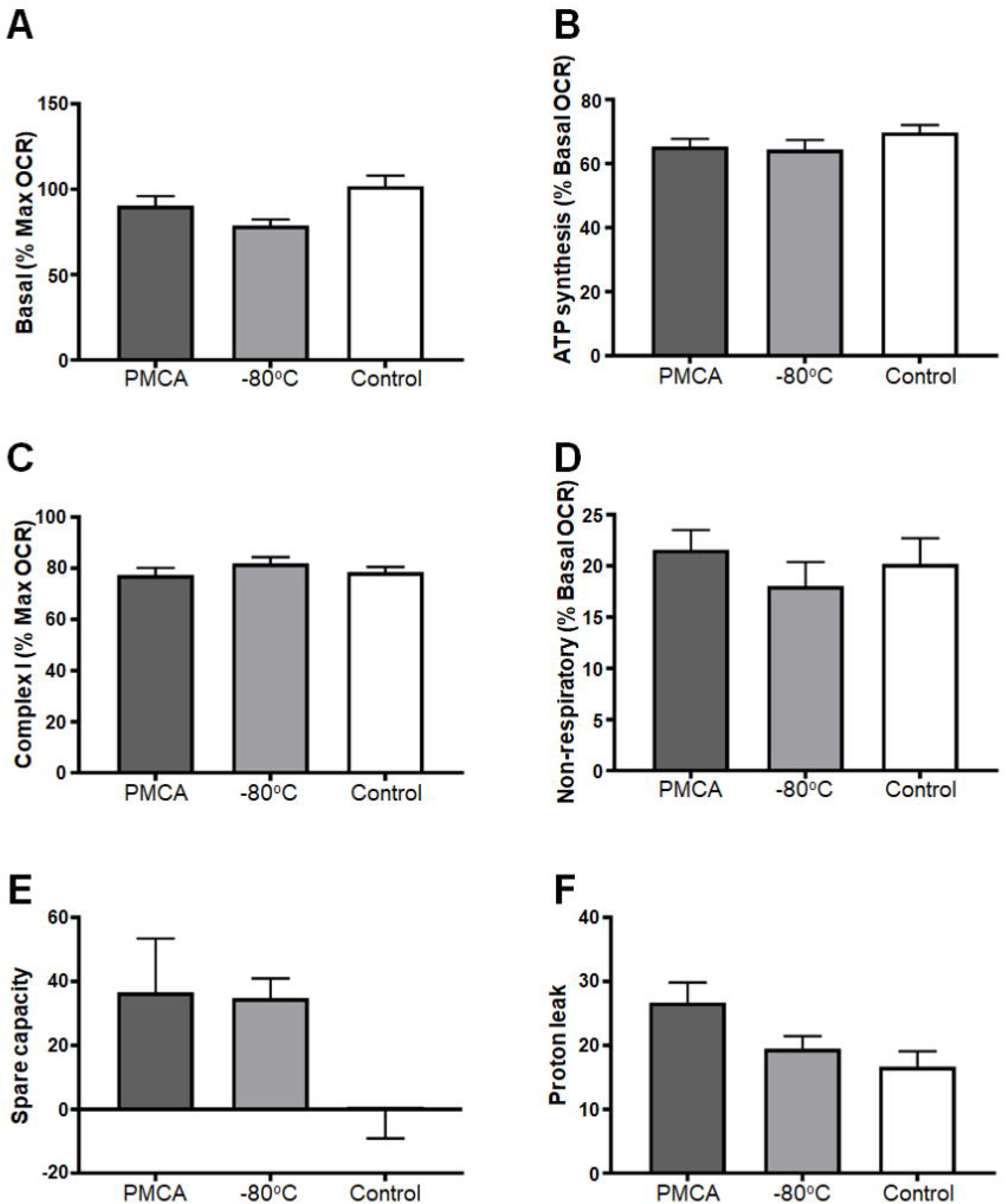
PMCA-generated misfolded α-synuclein do not cause functional deficits in mitochondria of SH-SY5Y cells. SH-SY5Y cells were incubated with PMCA-generated α-synuclein, monomeric α-synuclein (−80°C) or either buffer alone or left untreated which collectively represented the Control group. Media-containing cells were plated into a well of a Seahorse XFe24 plate pre-coated with Matrigel. The Seahorse XF Analyzer measured mitochondria respiration by detecting changes in oxygen (referred to as the oxygen consumption rate; OCR) following the addition of pharmacological agents: oligomycin, carbonyl cyanide m-chlorophenyl hydrazone (CCCP), rotenone and antimycin A. The functioning of mitochondria was determined as parameters expressed as % change max or basal OCR (depending on the functional readout). Spare capacity was measured as the difference between max and basal OCR and proton leak the difference between basal and ATP synthesis and non-respiratory. The average of five wells was taken for each sample per experimental replicate. Data presented as mean±SEM (n=16, 13, 18 for groups PMCA, −80°C and Control, respectively) and represent every experimental replicate performed. Statistical significance examined by ANOVA and Tukey’s multiple comparisons test with a statistical criterion of 0.01. No statistical significance was found for all experimental comparisons.

### Hyperactivity is not a result of altered glycolysis in SH-SY5Y neuroblastoma cells exposed to misfolded α-synuclein

Having identified the capacity of misfolded α-synuclein to cause hyperactive mitochondrial respiration, we next explored whether other components of cellular respiration were similarly affected. Glycolysis is an important upstream component to cellular respiration; it produces a small amount of the total ATP synthesized by the cell and critically produces acetyl co-enzyme A, which feeds into the tricarboxylic acid cycle, producing the electron donors FADH_2_ and NADH that are essential for oxidative phosphorylation. Given the intimate association between glycolysis and oxidative phosphorylation, the glycolytic potential of SH-SY5Y cells was assessed upon exposure to misfolded α-synuclein, which was measured via the extracellular acidification (ECAR) rate of cells in a glycolytic stress test. Within a cellular environment, ECAR is largely attributed to glycolysis and similar to the fluxes in cellular respiration, the glycolytic potential of cells may be tested by the sequential addition of compounds: glucose, oligomycin, rotenone & antimycin A and 2-deoxy-D glucose. The combination of these compounds, respectively, provide fuel for glycolysis to occur, inhibit ATP synthase, inhibit electron transport and inhibit glycolysis. The inhibition of electron transport by the combined actions of rotenone and antimycin A allows assessment of any contribution to ECAR by respiratory electron transport. Basic parameters calculated from this included starved ECAR, basal glycolytic ECAR, glycolytic capacity and spare glycolytic capacity. An image depicting a typical experiment measuring glycolytic potential is shown in Supplementary Figure 5A. Glycolytic potential of SH-SY5Y cells was measured upon exposure to PMCA-generated misfolded α-synuclein (PMCA), monomeric α-synuclein (−80°C), buffer alone (PBSN) and untreated (Untreat) with the same treatment regime as the mitochondrial respiratory experiments. An example experiment is shown in Supplementary Figure 5B and following three independent experiments, no difference was found between any of the experimental groups in all glycolytic parameters (Supplementary Figure 5C-F).

## Discussion

The PMCA is an assay shown to be capable of producing a range of various sized misfolded α-synuclein species (Herva et al., 2014). It was of interest to use this system to study the pathogenicity of misfolded α-synuclein, however further analysis on the species formed was limited due to a high concentration of detergent in the conversion buffer. Hence this study had two components: 1) to adapt the published protocol of PMCA-induced α-synuclein misfolding to enable their production to occur under non-toxic conditions and 2) to use this newly reformed system to investigate pathogenic mechanisms of misfolded α-synuclein. Replication of the original protocol was first achieved, with species produced via PMCA being characterized using a range of readouts (ThT, TEM and western immunoblot). This confirmed the system as a suitable tool to produce misfolded α-synuclein. The effectiveness of PMCA-induced α-synuclein misfolding when reconstituted in non-toxic buffers was next tested, and adaption of the protocol performed with the most efficient non-toxic buffer, PBSN. New parameters were determined to produce comparable misfolded content to the initial buffer, with 72 h being the optimized time to misfold α-synuclein in PBSN as assessed using a range of readouts. The ability of PBSN to be well tolerated in both normal, untransfected and transgenic PD-model neuroblastoma cells was the final assessment in the adaptation of the protocol and allowed further studies into the cellular targets of misfolded α-synuclein.

The major finding of this study was that PMCA-generated α-synuclein species affects mitochondrial respiration. Mitochondrial dysfunction is a central feature of PD and numerous studies have implicated α-synuclein in decreasing mitochondrial activity in various animal (Bender et al., 2013; Chinta et al., 2010; Li et al., 2013; Martin et al., 2006; Sarafian et al., 2013; Zhu et al., 2011) and cell culture models of disease (Devi et al., 2008; Parihar et al., 2008; Reeve et al., 2015). In addition to overall decreases in mitochondrial respiration, α-synuclein has been reported to cause functional deficits, particularly at the level of Complex I. Cells expressing wild-type or mutant α-synuclein reportedly have deficits in Complex I (Chinta et al., 2010; Devi et al., 2008), which are also observed in post mortem PD brain (Janetzky et al., 1994; Parker et al., 1989; Schapira et al., 1989). Hence the robust hyperactive respiration without functional deficit observed in SH-SY5Y cells in the presence of misfolded α-synuclein reported in the current study is in contrast to the general paradigm that α-synuclein decreases mitochondrial respiration in association with Complex I deficiency.

In order to delineate these divergent findings, it is important to consider factors that may contribute to variations in results. A major influence is likely to be the model used, with cell culture systems, transgenic animal models and human disease all harbouring very different cellular environments. Reporting an acute insult to naïve, healthy cells upon treatment with misfolded protein is likely dissimilar to the disease in humans which has been slowly progressing over years of clinical dormancy. Similarly, given that the overexpression of protein in cultured cells does not result in intracellular aggregation of protein, it also is unlike the human condition. In this regard, it could be argued the introduction of misfolded protein into naïve cells is more biologically relevant than a cellular environment where the clearance machinery is adequately removing protein aggregates. It is important to note that in the current study the delivery of α-synuclein into the cell was aided by lipofectamine, which could foreseeably have off-target effects associated with mitochondrial respiration. However, the inclusion of control samples treated with buffer only (PBSN) and lipofectamine showed equivalent readouts to cells untreated void of the transfection reagent. Cationic liposomal proteins are commonly used in studies delivering α-synuclein intracellularly (Luk et al., 2009; Nonaka et al., 2010) and we found its use did not contribute to mitochondrial readouts. Regardless, while each system identifies an effect of α-synuclein, its relevance to *in vivo* PD is uncertain. However, this also reflects an important point with data collected on human samples, which report the status of mitochondria at post mortem and hence terminal stages of disease. Detailing this stage of disease, while relevant, may not be an adequate representation of mitochondrial health for the majority of the disease process. Considering this, it is possible the mitochondria are able to adapt to the insult caused by α-synuclein for the majority of the disease course, before being overwhelmed at end-stage disease. Such a scenario would align with the findings of this study. In support of this, numerous studies on mitochondrial respiration in various cell types in small cohorts of ante mortem PD patients have failed to produce conclusive findings (Ambrosi et al., 2014; Barroso et al., 1993; Bravi et al., 1992; Cronin-Furman et al., 2013; Esteves et al., 2010; Martin et al., 1996; Parker and Swerdlow, 1998; Shinde and Pasupathy, 2006; Yoshino et al., 1992). Importantly, recently a larger study examining mitochondrial respiration in immortalized lymphocytes from 30 patients found enhanced respiratory activity with no functional deficit (Annesley et al., 2016). This elevation was found to be independent of age, disease duration or disease severity and hence supports the theory that mitochondria are adaptive in disease. Collectively, while further assessment is merited in models of disease, the data presented here using misfolded α-synuclein in non-transgenic cells suggests the dogma that mitochondria dysfunction is a persistent, overwhelming feature of PD may ultimately be revised.

Given that the PMCA produces a heterogeneous population of αsyn species, this work did not distinguish which species of misfolded αsyn is causing the changes to mitochondrial respiration. An attractive candidate is the oligomeric conformations given that these species have been shown to fragment CA-rich membranes (Nakamura et al., 2011), however, other species may be relevant. In transgenic mice overexpressing wild-type α-synuclein, the predominant species that accumulated in the mitochondria causing deficits was monomeric full-length N-terminally acetylated α-synuclein (Sarafian et al., 2013). The latter study highlights an important consideration to using recombinant protein to study pathogenic mechanisms of α-synuclein, which are not exposed to the numerous post-translational modifications endogenous α-synuclein is susceptible to. In PD, disease-associated α-synuclein is largely phosphorylated at position Ser129 (pSer129) (Fujiwara et al., 2002) and, in cultured cells expressing mutant A53T α-synuclein, both the stimulation of ROS production and alterations to mitochondrial respiration highly correlate with levels pSer129 (Perfeito et al., 2014). Therefore, the production of misfolded species using PMCA may not model all pathogenic subtypes of α-synuclein that present in a cell and cause damage in disease. An additional point is the buffer used in this study. High salt concentrations have been shown to modulate α-synuclein misfolding and hence further assessment will be necessary to examine the similarities between PMCA-generated α-synuclein produced in PBSN to protein misfolding that occurs *in vivo*.

This work did not elucidate whether the changes in mitochondrial respiratory function upon exposure to misfolded α-synuclein were due to a direct or indirect mechanism. The observation in Figure 3A that PMCA-generated α-synuclein species associates with CA is support for α-synuclein modulating mitochondria respiration via a direct interaction with this lipid. CA is commonly referred to as the signature phospholipid of the mitochondrion. It is predominately expressed in the IMM, which contains more than 40 fold higher levels of CA than the outer mitochondrial membrane (OMM) (de Kroon et al., 1997) and is a component of and/or required for the optimal functioning for all major components of the ETC (Paradies et al., 2014). Therefore, consistent with the elevated mitochondrial oxidative phosphorylation reported in this study, if an interaction between α-synuclein and CA occurs in a biological setting, it would likely result in broad spectrum changes to respiration. Further support for a direct interaction of α-synuclein with mitochondria comes from many studies showing mutant or wild-type α-synuclein associates with isolated mitochondria (Nakamura et al., 2008; Parihar et al., 2008) and mitochondria *in vivo* (Chinta et al., 2010; Cole et al., 2008; Devi et al., 2008; Parihar et al., 2008; Shavali et al., 2008). Also, α-synuclein has been shown to bind to membranes mimicking mitochondrial membranes, with a preference for those containing CA (Nakamura et al., 2011; Zigoneanu et al., 2012).

Given that this current work does not confirm a direct interaction between misfolded α-synuclein and mitochondria, it is possible that indirect mechanisms are driving the enhanced mitochondrial respiratory activity. As a general rule, enhanced respiration to all complexes without altering function would implicate an indirect mechanism. The most likely candidates for this are those that trigger an increase in ATP production. Autophagic flux is a major modulator of cellular respiration and thus the detection and clearance of misfolded α-synuclein may be a major contributing pathway to the changes in respiration seen here. However, the finding that glycolysis was unaffected upon exposure to PMCA-generated α-synuclein reveals important information on the potential mechanism of action. It suggests that if mitochondrial respiration was enhanced because of increased energy demands following exposure to misfolded α-synuclein, it did not also drive changes in glycolysis. This is consistent with the fact that no mitochondrial deficits were identified and hence glycolysis was not upregulated as a compensatory mechanism. It also eliminates changes to glycolysis as a driver of the effects seen. It should be borne in mind that both the glycolysis and mitochondrial function assays used here were designed to avoid constraints on activity by substrate supply rates. Furthermore, while further analysis is merited, observing hyperactive mitochondrial respiration following acute exposure of α-synuclein in this system is likely independent of chronic adaptive pathways.

In conclusion, the aim of this study was to develop a system to produce a heterogeneous population of misfolded α-synuclein species under non-toxic conditions and to investigate pathogenic mechanisms associated with these species. We show the development and characterization of a protocol to generate α-synuclein species in the non-toxic buffer PBSN using PMCA. These misfolded species were found to associate with the mitochondrial-specific lipid CA. CA accelerates α-synuclein fibrillization and alters the formation of α-synuclein species produced by PMCA. Critically PMCA-generated misfolded α-synuclein was shown to enhance mitochondrial respiratory activity in neuroblastoma SH-SY5Y cells. While it is important to note there are considerations with the work presented in the current study with regards to modelling disease-associated α-synuclein, the findings provide evidence that this mechanism can be a feature of misfolded α-synuclein and, hence, highlights the need for further assessment on the relationship between α-synuclein and mitochondria. The findings of this study provide important insight into pathogenic features of α-synuclein which may ultimately have implications for our understanding of the pathogenesis of synucleinopathies, including PD.

## Materials and methods

### Preparation of recombinant protein for α-synuclein PMCA

The open reading frame of the wild-type human α-synuclein gene SNCA had previously been cloned into the plasmid pRSET B under the control of the T7 promotor (Cappai et al., 2005). Recombinant human α-synuclein produced using this plasmid was purchased from Monash Protein Production facility (Monash University Clayton VIC) and supplied as 1 mL aliquots in distilled water (dH_2_O). Purchased protein was lyophilized overnight using a benchtop freeze dry system (Labcono, U.S.A) and stored at −80°C until reconstituted in buffer. The following buffers were used in this study: prion conversion buffer (PCB; 1% (v/v) Triton-X 100, 150 mM NaCl, cOmplete ULTRA protease inhibitor cocktail tablet (1 per 10 mL; Roche)/phosphate buffered saline (PBS; Amresco, Solon, OH, U.S.A)), PBS, PBS+150 mM NaCl (PBSN), tris buffered saline (TBS; Amresco, Solon, OH, U.S.A) and TBS+150 mM NaCl (TBSN). The final NaCl composition of buffers was 0.137 M, which increased to 0.287 M for those buffers with the additional 150 mM NaCl. For reconstitution of protein, 10 – 16 mg protein was dissolved in 1 mL buffer and centrifuged at 122, 500 *g* for 30 min, 4°C (Optima™ MAX Ultracentrifuge, Beckman Coulter) to sediment amorphous aggregates. Protein concentration of the supernatant was determined by measuring protein absorbance at 280 nm using a photometer (BioPhotometer, Eppendorf) and employing the Beer-Lambert law. Protein was diluted to a final concentration of 90 μM and 60 μL aliquoted into PCR tubes and stored at −80°C until experimental use in α-synuclein PMCA.

### α-synuclein PMCA

α-synuclein PMCA was performed as described previously (Herva et al., 2014). PCR tubes containing α-synuclein were briefly spun in a benchtop centrifuge (Microfuge®16 centrifuge, Beckman Coulter), 37±3 mg beads (1.0 mm zirconia/silica beads, BioSpec Products) were added and tubes sealed with Parafilm (Pechiney Plastic Packaging Company, Chicago, IL, USA). Samples were placed into a 96-tube rack in a microplate cup-horn sonicator (Misonix 4000) that was filled with water using a circulating water bath to maintain a steady state temperature of 37°C. PMCA reactions using α-synuclein consisted of 20 sec of sonication every 29 min 40 sec for varying lengths of time. Power setting 7 was used for all α-synuclein PMCA experiments. For protease resistance studies, PK was diluted in PBSN to 6 × the desired final concentration and 4 μL added to 20 μL of the PMCA product prior to incubation at 37 °C for 30 min with occasional agitation. Replacement of PK with PBSN in equivalent samples acted as non-PK controls and these samples were kept at room temperature (RT) during the digestion of PK-containing samples. All samples were then diluted in 4x NuPAGE LDS sample buffer (Life Technologies) and boiled at 100°C for 10 mins. Samples were resolved using SDS-PAGE and western immunoblot employed to detect α-synuclein species using the antibody MJFR1 (abcam138501, 1:10,000 dilution) (Brudek et al., 2016). To assess insoluble load of protein exposed to PMCA, ultracentrifugation was used. Following a PMCA reaction sample was diluted in PBSN to fill a 200 μL capacity thick-walled polyallomer tube (Beckman-Coulter). Samples were spun at 436,000 g for 1 h, room temperature. The pellet containing insoluble protein was washed in PBSN and centrifuged again under the same conditions. The supernatant was then discarded and the pellet resuspended in 8M urea/PBSN and incubated for 30 min, room temperature. Western immunoblot to detect human αsyn was performed as described previously.

### Thioflavin T (ThT) assay

Extent of fibrillization was quantified in PMCA products by reactivity to ThT (Sigma Aldrich). Here, protein solution was diluted 1/25 in ThT solution (20 μM ThT, 50 mM Glycine in dH_2_O, pH 8.5 with KOH) for a final volume of 250 μL in a well of a 96-well black polystyrene plate (Nunc™, Thermo Fisher Scientific). Fluorescence was measured using a Varioskan plate reader (Thermo Fischer Scientific). Triplicate wells of the same sample were averaged and values determined following subtraction of recordings of ThT solution void of sample to account for background fluorescence. When determining the effect of a compound on PMCA-induced α-synuclein fibrillization using ThT, separate tubes containing the compound in 60 μL of PBSN only was added to PMCA for 72 h, as well as protein containing the compound left at −80°C for the duration of the PMCA. These served as controls to confirm no ThT signal was attributed to the compound in the presence and absence of PMCA. For studies investigating kinetics of α-synuclein fibrillization, individual PCR tubes of α-synuclein were placed into the 96-tube rack of a running sonicator that was continuously cycling. This was done at specified intervals to enable all tubes to be removed at the same time such that their total process time corresponded to the time-point of interest.

### Transmission electron microscopy

For imaging PMCA products using TEM, samples were applied to formvar-coated grids and visualized by negative staining with uranyl acetate (2% w/v in H_2_O), with images captured using a Joel JEM-2100 electron microscope.

### Circular dichroism spectroscopy

Conformational changes in α-synuclein were investigated using circular dichroism spectroscopy essentially as described previously (Peverelli et al., 2016). Briefly, spectra were acquired using an Aviv Biomedical Model 420 CD spectrophotometer at 25 °C using 1 mm quartz cuvettes over a spectral range of 190–260 nm and using a step size of 0.5 nm, an averaging time of 4 s, and a slit bandwidth of 1 nm. Measurements for each sample or buffer were taken in triplicate, averaged, and reported as mean ± standard deviation. Prior to measurements, α-synuclein samples (in PBSN or PCB buffer) were diluted in ultrapure water to a final protein concentration of 100 μg L-1 or 65 μg L-1 for PMCA and PCB-treated samples, respectively. Readings were taken on α-synuclein protein either exposed to PMCA in PBSN (72 h), PCB (24 h) or left at −80 °C (0 h). Spectra for buffers alone, diluted equivalently in ultrapure water and averaged from triplicate measurements, was subtracted from protein spectra in all cases. Protein spectra are transformed into units of Δε (Kelly et al., 2005) to account for differences in protein concentration between samples.

### Analytical Ultracentrifugation

Analytical ultracentrifugation sedimentation velocity experiments were performed using a Beckman Coulter XL-A analytical ultracentrifuge equipped with absorption optics and using either a 4-hole An60 Ti or 8-hold An50 Ti rotor essentially as described previously (Gupta et al., 2018; Peverelli et al., 2016). Briefly, 2-channel epon centrepieces quartz cells were loaded with 300–380 μL of α-synuclein (initial protein concentration of 1.3 mg mL-1) and 320–400 μL of reference buffer (PBSN). Data were collected at either 25°C or 37°C continuously in absorbance mode using a wavelength of 232 nm, rotor speed of 42 000 rpm, radial distance of 6–7.3 cm, radial step size of 0.003 cm without averaging, and for up to 300 scans. An initial radial scan at a lower rotor speed of 3000 rpm was additionally collected for PMCA-generated α-synuclein samples. The partial specific volume of α-synuclein (0.7347 mL g-1), solvent density and viscosity were calculated as appropriate to the experiment temperature using the software SEDNTERP (available from http://www.jphilo.mail.com). Multiple time-staggered sedimentation velocity scans were fitted to c(M) distribution models using the software SEDFIT v16.1c (Schuck, 2000; Schuck et al., 2002). During fitting, the weight-average frictional ratio (f/f_0_), meniscus, and bottom radial positions were treated as floating parameters. Data were corrected for time-invariant noise, models solved to a resolution of 250, and maximum entropy regularization used (P = 0.68).

### Lipid binding assays

Lipid strips dotted with 15 long chain (>diC16) highly pure synthetic lipid analogues were purchased from Echelon Inc. and used to determine the affinity of PMCA-generated misfolded or −80°C monomeric (non-PMCA) α-synuclein to lipid species. Strips were blocked in 2% skim milk (2 h, RT), prior to incubation with 10 μg/mL α-synuclein protein diluted in blocking buffer for 2 h, RT. Extent of binding was measured using immunoblot by detection of α-synuclein using MJFR1 antibody.

### Preparation of lipids

500 μg of synthetic 16:0 cardiolipin (Echelon, Inc.) was diluted in a small volume of MeOH and solubilized by sonication (Misonix 3000) at 45 °C, 10 min. Cholesterol (Ajax Chemicals) was solubilised in a small volume of chloroform:MeOH (2:1), air dried and resuspended in MeOH. Lipid/MeOH was diluted to a final concentration of 741 or 74.1 μM and 5 μL added to PCR tubes containing 60 μL of 90 μM α-synuclein dissolved PBSN.

### Transfection of SH-SY5Y cells to produce stable cell lines overexpressing α-synuclein

In this study, all experiments with cultured cells used SH-SY5Y neuroblastoma cells that were confirmed mycoplasma negative, obtained from American Type Culture Collection (ATCC). To stably overexpress wild-type in SH-SY5Y, the α-synuclein:pcDNA3+ plasmid was used. SH-SY5Y cells were seeded in complete media (DMEM supplemented with 10% (v/v) fetal calf serum, 1% GlutaMAX (v/v) and 1% penicillin/streptomycin 100 × (v/v); Life Technologies) in a 12-well tissue culture plate at a density of 1.5 × 10^5^ for use 48 h later. Cells were serum starved in 750 μL serum free media for 4 h prior to transfection. Varying amounts of plasmid and lipofectamine transfection reagent (Thermo Fisher Scientific) were combined separately in 125 μL Opti-MEM (Thermo Fisher Scientific) per well, before being combined, mixed well and left to incubate for 20 min, RT. 250 μL of the combined solution was then added dropwise to each well and incubated in 37°C, 5% CO_2_ for 6 h before being replaced with fresh complete media. Cells were left to recover for 48 h before addition of the selective antibiotic G418 (300 μg/mL; Thermo Fisher Scientific) into complete media without added penicillin/streptomycin. Expression of α-synuclein in clones derived from transfected single cell colonies was monitored by western immunoblot analysis, with the cell line expressing the highest level of α-synuclein used for further experiments. The α-synuclein overexpressing SH-SY5Y cell line used in this study was derived from clone A4, and hence named as such.

### Lactate dehydrogenase (LDH) cytotoxicity assay

A lactate dehydrogenase (LDH) cytotoxicity assay kit (Thermo Fisher Scientific) was used to determine the cytotoxicity of buffers used in PMCA to A4 and untransfected SH-SY5Y cells. The protocol followed was as per the manufacturer’s instructions. Briefly, 1 × 10^4^ cells were seeded into a 96-well tissue culture plate and 24 h later triplicate wells were treated with 10 μL of PBSN or PCB diluted in dH_2_O to achieve a final concentration of 0.5, 1, 5, or 10% final volume of buffer. Additionally, one set of triplicate wells was treated with dH_2_O to act as a control for spontaneous cell death and one set was left untreated to be treated later with lysis buffer to measure maximum cell death (maximum LDH activity). Cells were incubated under standard cell culture conditions for 24 h. Following incubation, 10 μL lysis buffer was added to the set of untreated triplicate wells and incubated again for a further 45 min. 50 μL of media from all wells was then transferred to a 96-well flat-bottom plate. Reaction mixture (50 μL) was added to each well and incubated for 30 min RT protected from light followed by the addition of 50 μL of stop solution. Absorbance was measured at 490 nm and 680 nm to obtain sample fluorescence and background fluorescence, respectively. To determine LDH activity, the 680 nm absorbance was subtracted from the 490 nm value. Percent cytotoxicity of a sample was determined using the formula:

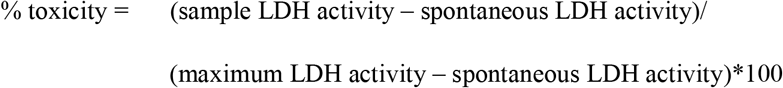

### Western immunoblot of cell lysates

All steps to lyse cultured cells were performed on ice using pre-chilled reagents. Pelleted cells were washed twice with cold PBS and resuspended in lysis buffer (150 mM NaCl, 50 mM Tris pH 7.4, 1% (v/v) Triton X-100, 1% (w/v) sodium deoxycholate (Sigma-Aldrich) and cOmplete ULTRA protease inhibitor in dH_2_O). Cells were left to lyse for 20 min with occasional agitation, then unwanted debris and nuclear material removed by centrifugation (1, 000 g, 3 min). The supernatant was collected and either used immediately in protein determination and western immunoblot analysis, or stored at −80°C for future use. Immunodetection of α-synuclein was performed using MJFR1 antibody.

### Detection of α-synuclein in transfected SH-SY5Y cells using immunofluorescence

3 × 10^4^ A4 and untransfected SH-SY5Y cells were seeded into 8-well Lab-Tek®II Chamber Slides (Nalge Nunc, Naperville, IL, U.S.A) and left for 48 h to strongly adhere. Cells were washed once in PBS and fixed with 4% (w/v) paraformaldehyde (PFA; Sigma-Aldrich) in PBS for 15 min, RT. Cells were washed three times in PBS to remove residual PFA and permeabilized in 0.5% (v/v) Triton X-100/PBS for 2 min, RT. Cells were washed again as described above and blocked in buffer (2% (w/v) bovine serum albumin (BSA; Sigma-Aldrich)/PBS) for 30 min, RT. Blocking buffer was removed and cells incubated with primary antibody (MJFR1 to detect α-synuclein) diluted in blocking buffer for 2 h, RT, followed by five washes in PBS. Fluorophore-conjugated (anti-rabbit, 568-Alexa Fluor conjugated) secondary antibody and 4’,6-diamidino-2-phenylindole (DAPI), also diluted in blocking buffer were then added to the chamber wells and left for 2 h, RT before a final five washes in PBS. Walls of the chamber slide were removed and mounted with glass cover slips (Mediglass, Taren Point, NSW, Australia) with ProLong™ Gold Antifade Mountant (Thermo-Fisher). Slides were inverted and left to dry overnight in the dark before being sealed with nail polish (Sally Hansen). Images were taken on a SP5 (Leica Microsystems Pty Ltd) confocal microscope.

### Preparation of samples for measuring cellular readouts using the Seahorse XFe24 Analyzer

When preparing α-synuclein and buffer control treatment samples for experiments using the Seahorse XFe24 Analyzer, autoclaved beads were used, and PBSN samples were generated by adding buffer (60 μL) to PCR tubes along with beads (equivalent to the setup with PMCA generated misfolded α-synuclein), and were similarly exposed to PMCA for 72 h. Confluent SH-SY5Y cells grown in T75 flasks were harvested, pelleted and resuspended in XF assay medium (unbuffered DMEM supplemented with 2.5 mM glucose and 1 mM sodium pyruvate). PMCA-generated misfolded and monomeric (−80°C) α-synuclein protein or PBSN only (26.25 μL per well) was mixed with lipofectamine (3 μL per well) and the combined solution added to the cells (2 × 10^5^ cells per well) in suspension and mixed gently. Equivalent cell densities were similarly left untreated. Cells and protein (525 μL in total) were then plated into the well of a Seahorse XFe24 plate that had been pre-coated (and subsequently air dried) with Matrigel (356231, Corning) diluted 1:2 in XF assay medium. Cells were left to adhere for 1 h at 37°C prior to analysis with the Seahorse XFe24 Analyzer with readouts calculated as per 200,000 cells.

### Measuring mitochondrial respiratory function in SH-SY5Y cells

Mitochondrial respiratory function in live SH-SY5Y cells was measured using the Seahorse XFe24 Extracellular Flux Analyzer (Seahorse Bioscience) via changes in the oxygen consumption rate (OCR) following the sequential addition of pharmacological agents (2 μM oligomycin, 1 μM CCCP, 5 μM rotenone, 1 μM antimycin A). Between each compound treatment, the average of three measurement cycles of OCR was taken, each cycle including a 3 min mix step, 2 min wait and a measurement time of 3 min. Each condition tested had a minimum of three replicate wells, with the average of the values taken per experiment. Equipment setup for glycolysis measurements was as per respiratory function experiments and glycolytic function determined from changes in pH associated with the extracellular acidification rate (ECAR) in live SH-SY5Y cells. This was measured following the successive addition of pharmacological agents (10 mM glucose, 2 μM oligomycin, 5 μM rotenone 5 μM & antimycin A, 100 mM 2-deoxy-D glucose).

## Data analysis

Densitometric analysis was performed on unsaturated western immunoblot images using Image Lab. All statistical analysis was performed using GraphPad Prism. A statistical criterion of 0.05 was used for all experiments except for mitochondrial respiration data which used 0.01. Normality was assessed on all samples subjected to statistical analysis to ensure data met the assumptions of the tests used and statistical outliers identified. When values from at least three independent replicates were combined, they were depicted as mean±standard error of the mean (SEM).

## Acknowledgements

The authors wish to acknowledge Peter Lock and Ben Scicluna for technical contributions to this manuscript. Roberto Cappai kindly supplied both α-synuclein constructs (α-synuclein:pcDNA3+ and α-synuclein:pRSET B) used in the project. The TEM images were obtained with the help of the Bioimaging Platform (La Trobe University) and the Biological Optical Microscopy Platform (The University of Melbourne). Acknowledge also goes to the La Trobe University-Comprehensive Proteomics Platform for providing infrastructure and expertise.

