## Supplementary Data for "Misfolded α-synuclein causes hyperactive respiration without functional deficit in live neuroblastoma cells"

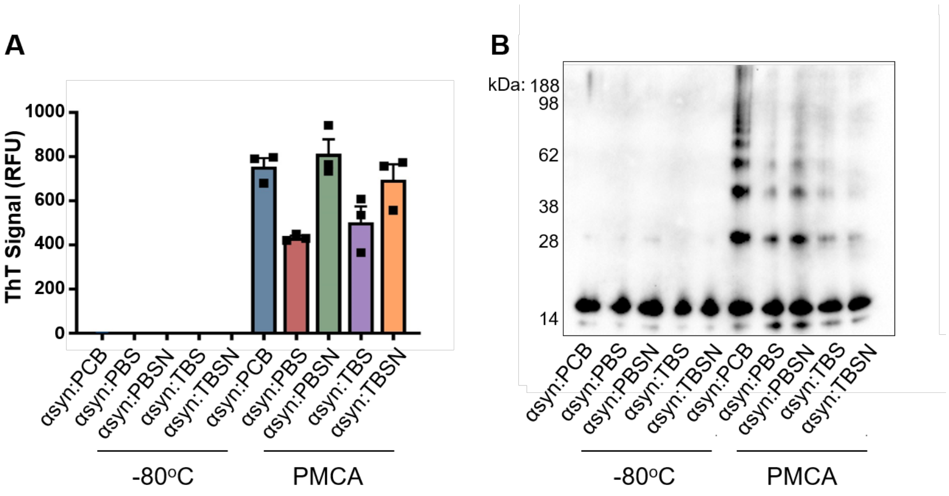


**Supplementary Figure 1: PMCA-induced α-synuclein fibrillization can occur under non-toxic conditions.**

Lyophilized recombinant wild-type protein was reconstituted in PCB, PBS, PBSN, TBS or TBSN and subjected to PMCA for a total process time of 24 h. Equivalent samples not subjected to PMCA (-80ºC) served as the non-PMCA monomeric controls. Extent of fibrillization in PMCA-exposed samples was assessed using (A) ThT fluorescence, where the values obtained for each experimental replicate represented the average fluorescence of triplicate wells after subtraction of a blank well to account for background fluorescence (RFU= relative fluorescent units). Data presented as mean±SEM (n=3) and represent every experimental replicate performed. (B) Western immunoblot analysis using α-synuclein-specific monoclonal antibody MJFR1 (amino acid specificity: 118-123) n=1.

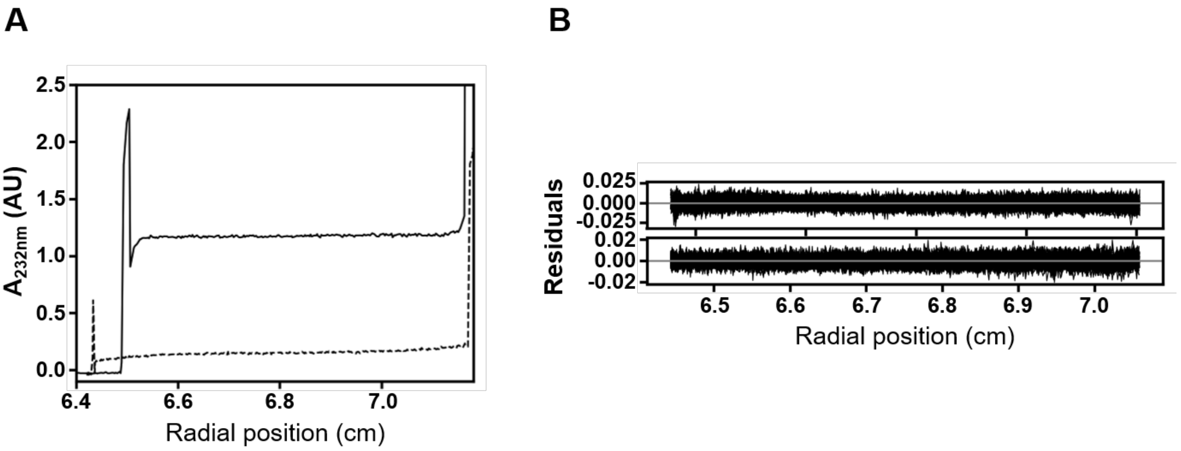


**Supplementary Figure 2: Analytical ultracentrifugation parameters.**

Absorbance (232 nm) plotted as a function of radial distance for 72 h PMCA α-synuclein:PBSN at rotor speeds of 3000 rpm and 42,000 rpm (solid and broken lines, respectively). Initial scans at each speed are represented. (B) Residuals for fits of a-synuclein analytical ultracentrifugation sedimentation velocity data to the c(M) distribution model. The residual value of 0 is indicated by a solid grey line. Top: 0 h a-synuclein:PBSN, bottom: 72 h a-synuclein:PBSN.

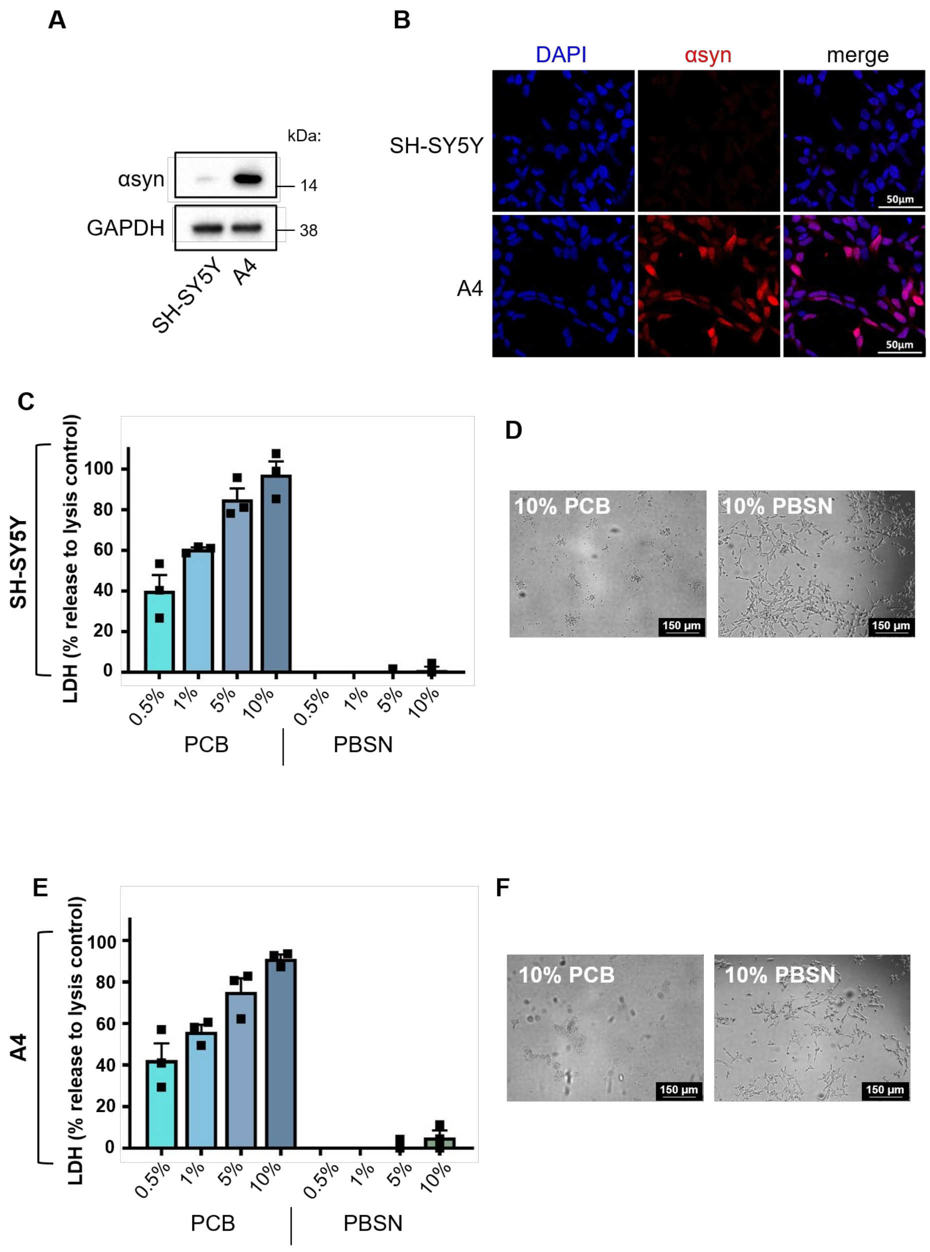


***Supplementary Figure 3: PBSN buffer is non-toxic to α-synuclein overexpressing SH-SY5Y cells (A4).***

SH-SY5Y cells were stably transfected with human wild-type α-synuclein. The single cell colony, A4, was shown to express elevated levels of α-synuclein compared to untransfected SH-SY5Y cells via (A) western immunoblot and (B) immunofluorescence using α-synuclein-specific monoclonal antibody MJFR1 (amino acid specificity: 118-123). *(C,E)* Toxicity of PCB and PBSN in untransfected Sh-SY5Y and A4 cells after 24 h was measured by assessing LDH levels via absorbance. Values were expressed as a percent cytotoxicity compared to equivalent cells treated with lysis buffer (maximum cell death). Deduction of water treated samples accounted for spontaneous LDH release. All buffers were measured using triplicate reads. Data presented as mean±SEM (n=3) and represent every experimental replicate performed. (D,F) Following treatment with PBSN or PCB DIC images of cells were taken. n=1, representative images.

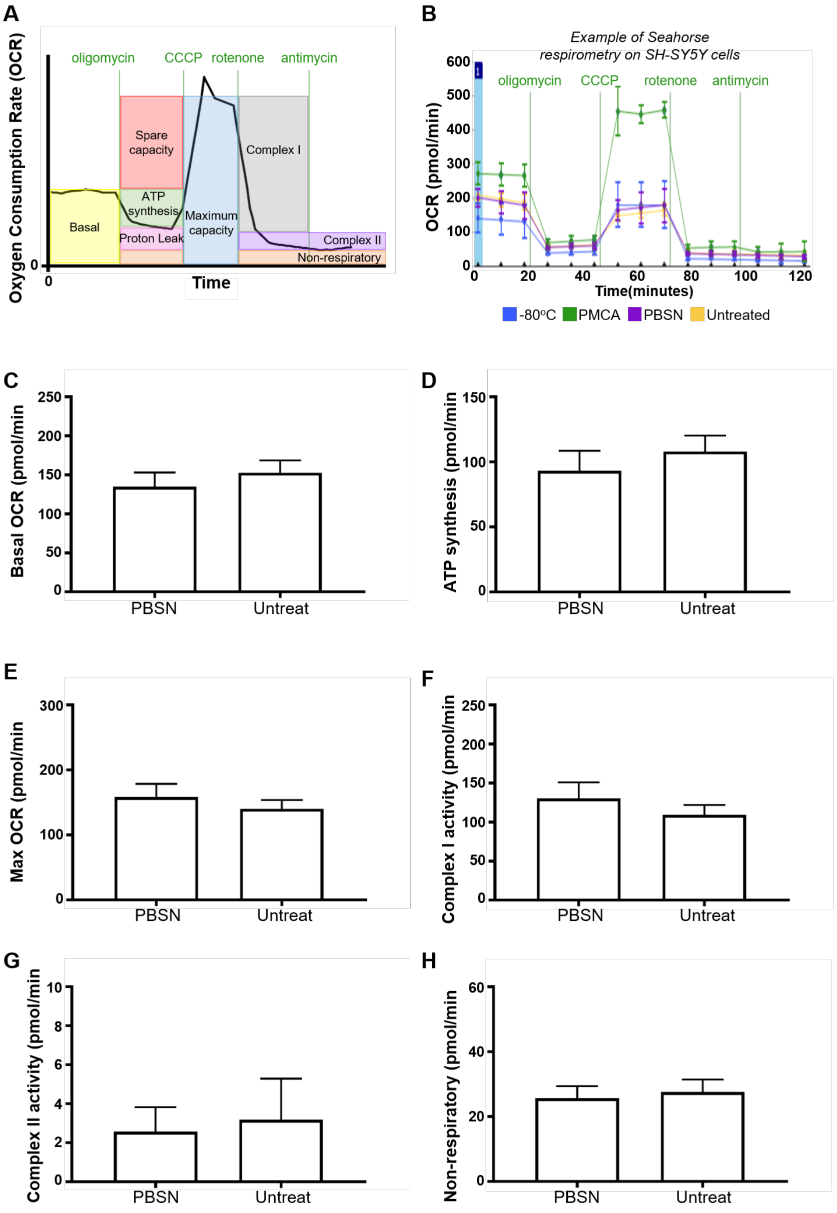


**Supplementary Figure 4: Mitochondrial respiration in SH-SY5Y cells treated with PBSN or untreated.**

SH-SY5Y cells were incubated with PBSN or left untreated (Untreat). Media-containing cells were plated into a well of a Seahorse XFe24 plate and mitochondria respiration measured in adhered cells using the Seahorse XF Analyzer. This was performed by detecting changes in oxygen (referred to as the oxygen consumption rate; OCR) following the addition of pharmacological agents: oligomycin, carbonyl cyanide m-chlorophenyl hydrazone (CCCP), rotenone and antimycin A. (A) A schematic of the respiration readouts measured and (B) raw OCR values obtained during the experiment. The following parameters were measured: (C) basal OCR, (D) ATP synthesis, (E) max OCR, (F) complex I activity, (G) complex II activity and (H) non-respiratory. For each experimental replicate, the average of multiple wells was taken for every sample. Data presented as mean±SEM (n=6, 12 for groups PBSN and Untreat, respectively). Statistical significance was examined by Student's t-test with a statistical criterion of 0.05.

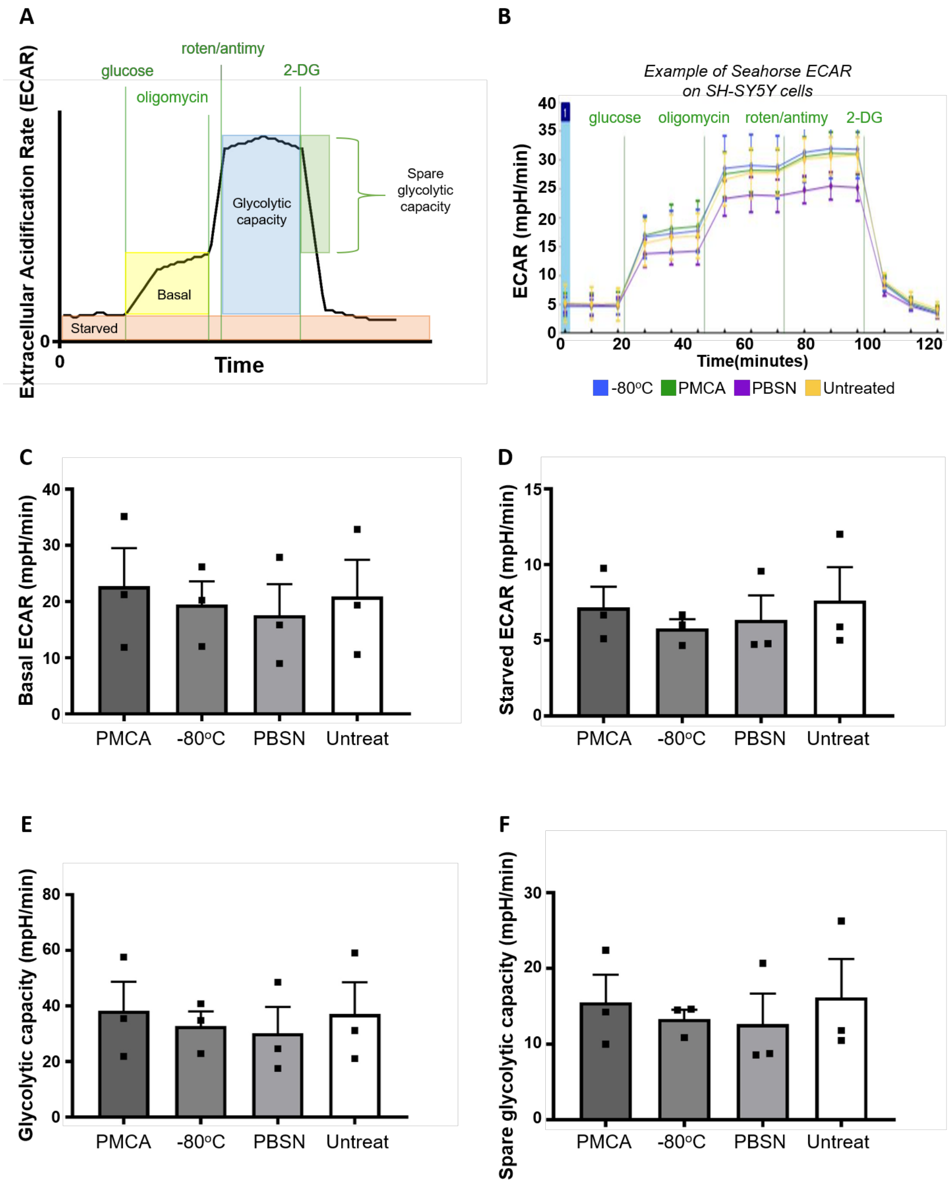


***Supplementary Figure 5: PMCA-generated misfolded α-synuclein do not affect glycolysis in SH-SY5Y cells.***

SH-SY5Y cells were incubated with PMCA-generated α-synuclein, monomeric α-synuclein (-80ºC), buffer alone (PBSN) or left untreated (Untreat). Media-containing cells were plated into a well of a Seahorse XFe24 plate pre-coated with Matrigel. The Seahorse XF Analyzer measured glycolytic potential by detecting changes in pH (that corresponds to the extracellular acidification rate of a cell; ECAR) in the absence and presence of glucose, and following the addition of pharmacological agents: oligomycin, rotenone+antimycin A (roten/antimy) and 2-deoxy-D glucose (2-DG). In doing so, the following parameters were measured: basal ECAR (A), starved ECAR (B), glycolytic capacity (C), spare glycolytic capacity (D). The average of five wells was taken for each sample per experimental replicate. Data presented as mean±SEM (n=3, all groups) and represent every experimental replicate performed. Statistical significance examined by ANOVA and Tukey's multiple comparisons test with a statistical criterion of 0.05. No statistical significance was found for all experimental comparisons.

| | Comparison | Values ± SEM (pmol/min) | p-value | Significance |
|---|---|---|---|---|
| **Basal** | PMCA vs. -80$^{o}$C | 202.2±13.66 and 113.8±7.87 | <0.0001 | **** |
| | PMCA vs. Control | 202.2±13.66 and 146.8±12.07 | 0.0022 | ** |
| **ATP Synthesis** | PMCA vs. -80$^{o}$C | 131.1±8.95 and 72.90±5.20 | 0.0001 | *** |
| **Max** | PMCA vs. -80$^{o}$C | 238.9±24.27 and 148.6±12.23 | 0.0025 | ** |
| | PMCA vs. Control | 238.9±24.27 and 146.5±11.07 | 0.0008 | *** |
| **Complex I** | PMCA vs. Control | 188.1±23.18 and 116.5±10.63 | 0.0062 | ** |
| **Non-respiratory** | PMCA vs. -80$^{o}$C | 44.41±5.88 and 21.38±3.50 | 0.0013 | ** |
| | PMCA vs. Control | 44.41±5.88 and 27.00±2.75 | 0.0091 | ** |

**Supplementary Table 1: Values of significant differences identified in Figure 4.**

Statistical significance was examined by ANOVA and Tukey's multiple comparisons test with a statistical criterion of 0.05. *p<0.05, **p<0.01
